# A network based approach to identifying correlations between phylogeny, morphological traits and occurrence information of fish Species in US river basins

**DOI:** 10.1101/2023.01.09.523236

**Authors:** Richa Tripathi, Amit Reza, Guohuan Su, Adam Mertel, Justin M. Calabrese

## Abstract

The complex network framework has been successfully used to model interactions between entities in Complex Systems in the Biological Sciences such as Proteomics, Genomics, Neuroscience, and Ecology. Networks of organisms at different spatial scales and in different ecosystems have provided insights into community assembly patterns and emergent properties of ecological systems. In the present work, we investigate two questions pertaining to fish species assembly rules in US river basins, a) if morphologically similar fish species also tend to be phylogenetically closer, and b) to what extent are co-occurring species that are phylogentically close also morphologically similar? For the first question, we construct a network of Hydrologic Unit Code 8 (HUC8) regions as nodes with interaction strengths (edges) governed by the number of common species. For each of the modules of this network, which are found to be geographically separated, there is differential yet significant evidence that phylogenetic distance predicts morphological distance. For the second question, we construct and analyze nearest neighbor directed networks of species based on their morphological distances and phylogenetic distances. Through module detection on these networks and comparing the module-level mean phylogenetic distance and mean morphological distance with the number of basins of common occurrence of species in modules, we find that both phylogeny and morphology of species have significant roles in governing species co-occurrence, i.e. phylogenetically and morphologically distant species tend to co-exist more. In addition, between the two quantities (morphological distance and phylogentic distance), we find that morphological distance is a stronger determinant of species co-occurrences.

## I. INTRODUCTION

Functional traits and phylogenetic relatedness are important attributes of species in a community to assess the two critical facets of biodiversity i.e., functional and phylogenetic diversity. It is often assumed that the functional diversity of animal and plant species are concordant with their phylogenetic diversity [1–3]. Furthermore, for optimizing conservation strategies, it is crucial to understand whether conserving phylogenetic diversity will help in conserving functional diversity of species [4–6]. The relationship between functional traits and phylogenetic closeness has been widely explored in various taxa in the past decades. Recently, there have been many studies to better understand whether ecologically relevant functional traits are conserved along the phylogeny. To do this, *phylogenetic signal* is typically addressed by testing if closely related species share more similar traits than expected by chance [7–9].

The functional and phylogenetic similarity between co-occurring species also plays a key role in the processes governing community assembly [10, 11]. Understanding the determinants of species co-occurrence in communities is a fundamental question in ecology [12]; therefore a few studies [11, 13] have focused on the structure of communities in species interaction networks as a way of gaining insights on coexistence mechanisms. There are two mutually exclusive mechanisms to illustrate the similarity of co-occurring species in a community. The competition mechanism [14–16] predicts that if a community is structured by competition, then species within the community should be more different functionally and phylogenetically than expected from assemblages that are comprised of random samples drawn from the potential species pool. In contrast, the environmental filtering mechanism predicts that if a particular type of habitat requires possession of certain adaptive traits, species within the communities should have more similar traits or be more closely related than expected by chance [17]. Ultimately, patterns of species co-occurrence may depend on how both biotic interactions and environmental filtering act over ecological and evolutionary time scales [18]. For instance, Winston (2005) [19] reported that functionally similar species co-occurred less frequently than more dissimilar species for 219 fish communities from the Red River basin in the US, supporting the biotic interactions mechanism. In contrast, Peres-Neto (2004) [17] reported that co-occurring species had greater functional and phylogenetic similarity than expected by chance for stream fish species in the Macau River basin in Brazil, and concluded that the co-occurrence patterns were mainly driven by the environmental filtering. As the organization of species into communities is a non-random process constrained by their interactions and system dynamics, it is important to identify such community assembly rules against the backdrop of species’ evolutionary history. Hence, a framework is needed that incorporates, phylogenetic and morphological relatedness of species, to reveal their groups (communities). Further, using statistical properties of these groups, one can test if these species also tend to co-occur. Hence, such an analysis can help understand if and by what degree does morphological and phylogenetic closeness impacts co-occurrence of species and if they have a differential impact on community assembly rules.

In this work, we complied occurrence data of native freshwater fishes from 2073 watersheds in the US. We measured 10 morphological traits that related to fish functions [20] and subsequently compiled phylogenetic information on these fish species [21]. We test the relatedness of species phylogeny with species morphology; and their relation with the species co-occurrences. For this, we used the concept of complex networks. The complex network framework not only allows the modeling of one-to-one interactions (relatedness) between entities of a complex system, but can also explain the emergent structural patterns that have implications for its function. Ecological Networks are finding increasing applicability in the study of species interactions, spatial ecology, community assembly, and its relation with phylogeny [22–24]. Studies have used fish co-occurrence networks for identifying a spatial cluster of fish-subgroups [25], and phylogenetic networks to retrace species dispersal history [26] in freshwater fishes in Ontario, Canada. While most of these network studies deal with species interactions based on only one of their attributes (e.g., phylogeny, or morphology, or co-occurrence), network studies accounting for multiple attributes of species that might govern their interactions are not very common. Therefore in the present network based study, we specifically 1) test whether morphological closeness of fish species is related to phylogenetic closeness (for HUC8 level regions of high fish co-occurrence), and how this relation changes geographically, and 2) explore whether trait similarity of fish species and their phylogenetic closeness predicts their co-occurrence in basins, and to what degree. The two classes of networks we constructed from the datasets are 1) co-occurrence based networks of HUC8 regions, and 2) nearest-neighbour directed networks of species based on the metrics of phylogenetic distances and morphological distances to other species in the system. Through module-extraction on these networks, which identifies groups of densely connected entities in the network, we quantify relationships among these quantities. In summary, our goal is to design a multi-attribute network-based analysis, where phylogenetic and morphological relatedness, and co-occurrence between fish species are simultaneously taken into account so as to be able to understand correlations between them.

## II. ABOUT THE DATASET

### 1. Species occurrence data

We compiled native species occurrence records at the watershed scale (i.e., Hydrologic Unit Code 8; HUC8) from NatureServe (https://www.natureserve.org) and included both extant and extinct species to account for species historically present in a given watershed. Records of non-native species occurrence were excluded from the entire study. Overall, the occurrence dataset has presence and absence information of 804 fish species in 2073 HUC8 regions.

### 2. Functional traits

We compiled ten morphological traits related to fish locomotion and food acquisition from the FISHMORPH database [27, 28], and FishBase [29]. The ten morphological traits include maximum body length (Length), body elongation (BlBd), relative eye size (EdHd), oral gap position (MoBd), relative maxillary length (JlHd), vertical eye position (EhBd), body laterally shape (HdBd), pectoral fin vertical position (PFlBl), pectoral fin size (PFiBd), and caudal peduncle throttling (CFdCP). Due to insufficient information on some species, some values were missing in the raw functional trait data. We statistically imputed these missing values [20] with a machine learning algorithm called ‘missForest’ [30, 31]. This method uses a random forest trained on the observed values of a data matrix to predict the missing values and automatically calibrates the filling values by a set of iterations [32].

### 3. Phylogenetic relatedness

We obtained the phylogenetic information of these fish species from Rabosky et al. (2018) [21], which includes 31, 526 marine and freshwater ray-finned fishes. This dataset is based on 11, 638 species whose position was estimated from genetic data; the remaining 19, 888 species were placed in the tree using stochastic polytomy resolution. We pruned the tree and kept only the 804 native freshwater fish species used in our study. Then we computed the pairwise distances between the pairs of these species from the pruned phylogenetic tree by using the ‘cophenetic.phylo’ function from the R package “ape”[33].

## III. METHODOLOGY

We use the phylogenetic distance (PD) dataset to measure phylogenetic closeness and farness between the fish species. In the trait dataset, 10 morphological traits of all species are normalized (by division by maximum value of the trait) to avoid the impact of their magnitudes on the analysis. Further, we assume them as vectors in a 10-dimensional trait space, and visualize each species as points in this vector space. For assessing the trait resemblances of species from the morphological traits dataset, we needed to define a measure and therefore we borrow the concept of cosine-similarity (CS). Hence, for measuring the trait-similarity between species, we use the CS [34–36] measure. The CS between trait vectors is the angular distance (*θ*) between the species in the 10-dimensional feature space, measured as the cosine of *θ*. Hence, unlike the Euclidean distance, it is independent of the vector magnitudes, with CS = 1 implying that the vectors are co-linear or that the species are similar and CS = 0 that the vectors are orthogonal or that the species are not similar in their traits. However, because we intend to correlate trait similarity with phylogeny, which is conventionally expressed as phylogenetic *distance* (PD), we use cosine-distance (CD) defined as CD = 1 – CS, instead of CS, and call it morphological distance between species throughout our analyses for consistency.

### A. Complex Network framework

First category of networks that we study is the basin network. In the basin network, the nodes are 2073 HUC8 regions and edges depict the number of fish species these basins share. Hence, absence of an edge would depict no common species between the basins. This network is weighted and un-directed by nature, where edge-weights are the number of species the two basins in question, share between them. We perform module-detection on the basin network as well to identify groups of basins with higher co-occurrences and their geographical distribution. Further, to infer dependence between CD and PD, we look at basin-level mean CD (〈CD〉) and mean PD (〈CD〉) over species in the basin and obtain the linear association between them.

Studying species interactions in a network framework allows us to gauge the overall connection structure in a single snapshot. As a second category of networks, we construct species networks wherein nodes (species) form the edges/ links to other nodes based on the PD or CD between them. If the two species are phylogenetically closer to each other, i.e. they have small PD between them, they are connected via an edge, and similarly, other node pairs in the network form their connections. The number of connections beginning from all source nodes are fixed (NN) beforehand and each of them have NN outgoing connections. This results in directed networks wherein edges capture species-specific interactions or the asymmetry of interactions. For example, if species P is among the top nearest phylogentic neighbours of species Q, it does not imply that species Q is among the top nearest phylogentic neighbours of species P. Hence, if a directed edge exist from Q to P, it may not exist from P to Q. In this manner, we identify a fixed number of nearest neighbors (NN) nodes for each node in the network and obtain directed networks of fish species. The procedure for constructing species networks based on a pre-defined number of NN is explained in Figure. 1. The matrix defining the connections in the network is called the adjacency matrix (**A**), where *A_ij_* = 1 denotes the presence of connection and *A_ij_* = 0 denotes the absence of the connection between nodes **i** and **j**. As described, these are directed networks (identified by the presence of directed links between the nodes as shown in Figure. 1), as each of the species selects its NNs depending on the value of its PD or CD to target species. The two kinds of species’ networks, constructed and analyzed in this work are:

- PD based Network: Each species in the network connects to *n* other species that are phylogenetically closest to itself.
- CD based Network: Each species connects to n other species that are trait-wise most similar to itself.

**FIG. 1.**
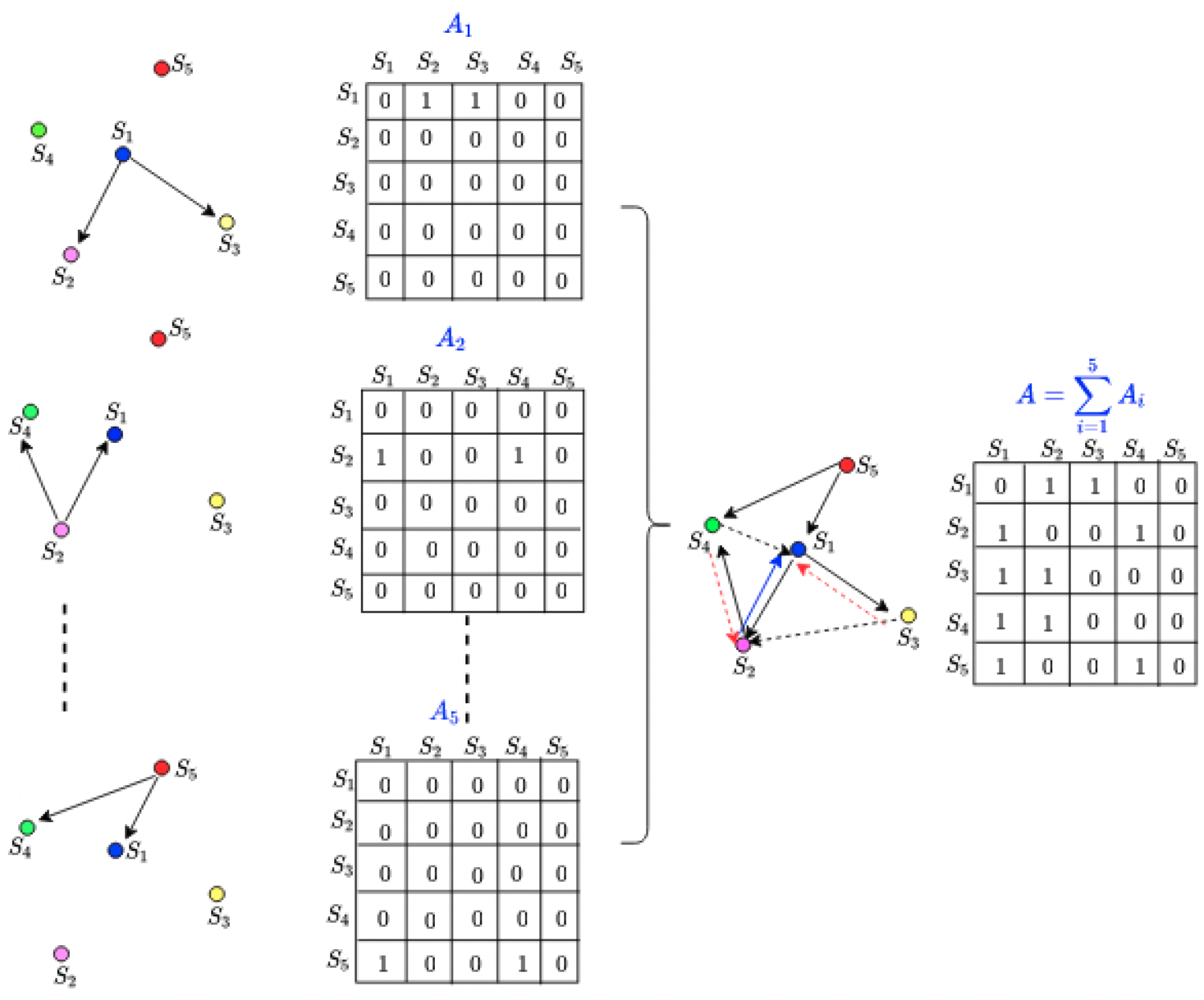
Procedure for construction of network of species. The schematic diagram demonstrates the procedure for construction of network of species based on their NNs. The panel on the right shows an example network of five species, with each node connecting to two of its NNs. This network (main) is formed by aggregation of different sub-networks. The panel on the left shows these sub-networks; first sub-network shows species *S*_1_ connecting to *S*_2_ and *S*_3_, second shows *S*_2_ connecting to *S*_1_ and *S*_4_, and so on. Alongside each of these sub-networks and the main network is shown an adjacency matrix, where 1 (0) in a cell indicated the presence (absence) of connection between the nodes. The network on the right shows all the connections in the sub-networks on the left, and its adjacency matrix is the sum of all the sub-network adjacency matrices. The solid edges are the ones explicitly shown in the sub-networks, and the dashed edges are edges from sub-networks whose display has been skipped on the left.

Hence, for PD based species network, we connect all the 804 species to their *n* phylogenetic nearest neighbours, and for CD based species network, we connect them to their *n* trait-wise nearest neighbours. Further, for both categories of these species networks we identify modules (or groups) of species using a network community-detection algorithm and then explore co-occurrence of the species within modules in the HUC8 regions. For species within each of these modules, we obtain mean over pair-wise PD (〈PD〉) and mean over pair-wise CD (〈CD〉). Following this, we obtain the number of HUC8 regions in which at least two of the species within the clusters occur together. Hence, each of the species clusters is assigned three quantities: 〈CD〉, 〈PD〉 and the number of basins of co-occurrence of at least two species. We denote the last quantity as NBS in the upcoming text.

As stated earlier, our goal is to ascertain which of these metrics, actually and significantly plays a role in community assembly rules. Hence, we define two null hypotheses: *H*_1_ and *H*_2_ as follows.

1. *H*_1_: The number of basins of co-occurrence of species are not related to their trait similarity.
2. *H*_2_: The number of basins of co-occurrence of species are not related to their phylogenetic closeness.

Notice that the averages (denoted by 〈x〉) in the species network depict the mean of pair-wise x over species in a given module, whereas in the basin network they are the mean of pair-wise x over species in a given basin (HUC8). To summarize, we construct and analyze both of these kinds of networks (basin network and species networks) to assess the co-dependence between the three quantities, i.e. phylogenetic distance, morphological distance and probability of co-occurrence of species. While the basin network is helpful in understanding the geographical distributions of phylogenetic and morphological diversity, the species networks help us find meaningful species communities, their collective chances of occurrences vis-à-vis their phylogentic and morphological relation. The following section explains the significance of and method for identifying modules in the networks.

### B. Communities in the network

An important concept in a network study is that of a module [37]. A module is a set of nodes that are more densely connected among themselves than to the rest of the network. For example, in a collaboration network, this would mean a group of authors that write papers with each other more frequently than with other authors in the network. For our basin network, a module is a subset of basins that on average share more species within the subset than with the remaining basins. The identification and visualization of these modules can help in understanding which basins tend to cluster together and if the modules are restricted to specific geographical locations on the network. Similarly, a module in a network of species is a group of species that are phylogenetically (if the network is constructed based on PD) or trait-wise (if the network is constructed based on CD) closer to each other, than to the rest of species. To identify modules in the undirected basin network, we use the Louvain algorithm [38] that returns the best partition of the network into modules by optimizing network modularity (Q) [39]. For the species networks, we use Leiden algorithm [40], that also used modularity and can efficiently identify modules in directed networks. A high value of *Q* (close to 1) indicates high divisibility of the network into clearly defined modules and vice-versa. Therefore, module compositions of the network are robust representations of the clustering of network nodes when the modularity is high, i.e., close to 1.

## IV. RESULTS

Here we present our results from the analysis of basin network first and then from the species networks.

### A. Relation between species phylogenetic distances and trait dissimilarity at the watershed level: Analysis of basin network

Using the basin network, we explore the relation between mean phylogenetic distances and mean trait dissimilarity of species within basins in the context of their geographical occurrence. Due to a higher species richness in the eastern part of the US than the western part (see Figure. 5 (c) in the supplementary information (SI)), the network is denser in the eastern part, signifying a larger number of co-occurrences of species in those basins. On performing the module detection on the basin network, geographically clustered groups of basins are obtained. By definition, basins within a cluster share more species among them than they share with basins in other clusters. The obtained communities are shown in different colours on the US map in Figure. 2 (a) (the network edges are lightened for visualization purposes). It is clear from the map that, as expected, the geographically close regions have larger co-occurrences of species than those that are farther apart. Additionally the clusters on the West end (red) and East end (orange) of the map seem to be broadly demarcated from their adjacent clusters by geographical divides that separate watersheds flowing into different oceans (shown by black bold lines on the map).

**FIG. 2.**
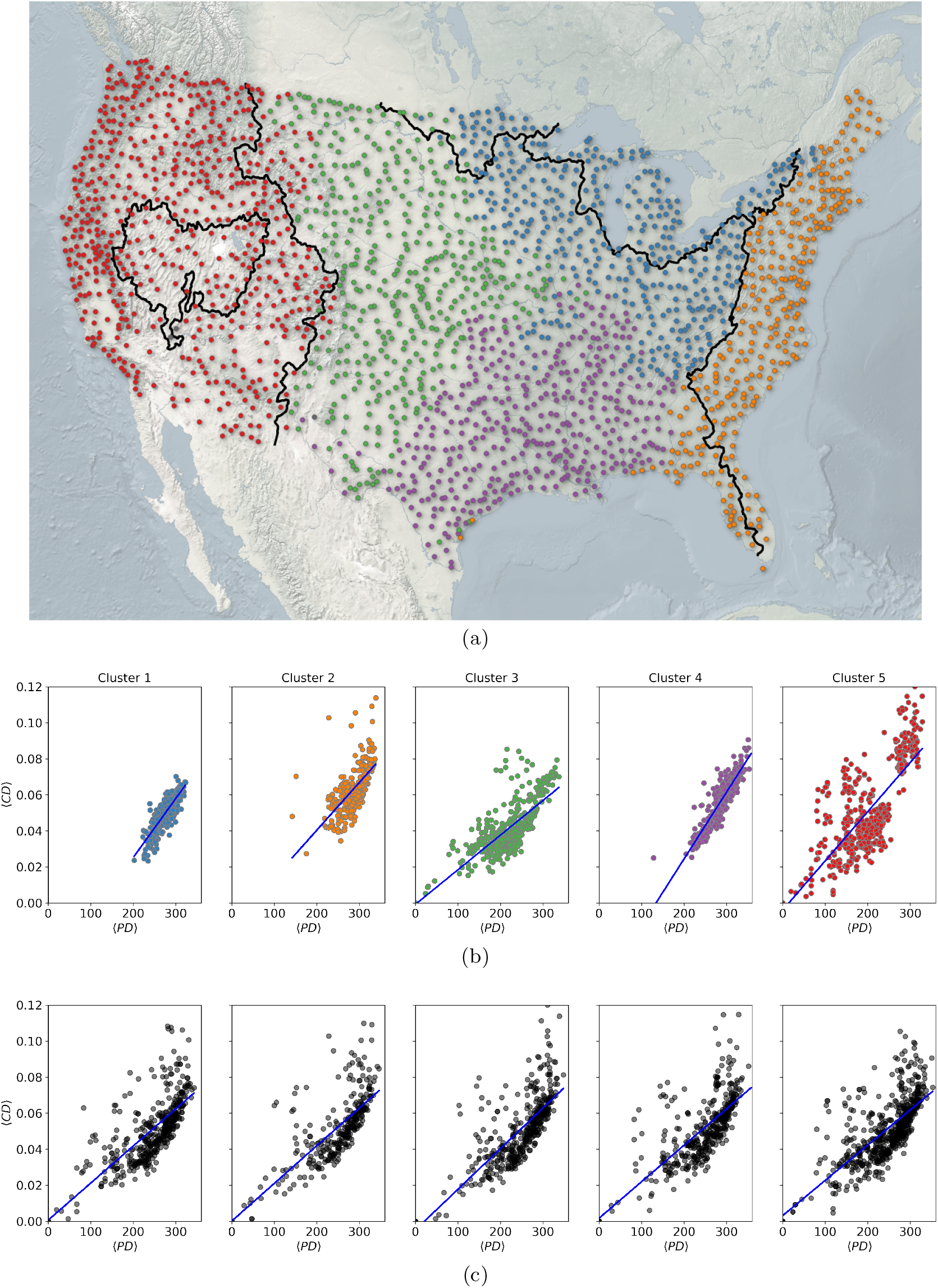
Identification of clusters of HUC8s based on number of species of co-occurrence. The figure shows a network of US watersheds with nodes plotted at the centroids of 2073 HUCs and colored according to the cluster they belong to, the edges are lightened for visualization. Basins in only the coterminous United States is shown here. The clusters are the modules identified using community detection algorithm on the network of HUCs, where the network is constructed based on number common species between HUCs. (b) The figure shows 〈CD〉 versus 〈PD〉 of the species in the basins in the HUC8 clusters. The colors are indicative of the module index. (c) The figure shows 〈CD〉 versus 〈PD〉 of randomly selected basins from across the network. The number of randomly selected basins are same as in the actual modules shown in part (b). The red lines in parts (b) and (c) are the best linear fits to the data.

For correlating co-occurrence information with phylogenetic closeness and trait similarity, we obtain basin-wise means of PD and CD of species within each basin. On plotting mean CD (〈CD〉) versus mean PD (〈PD〉), for each of these five basin clusters, we make two interesting observations. The first one supports the correlation between 〈CD〉 and 〈PD〉 of the species as can be seen from subplots in Figure. 2 (b) for all the five clusters. We perform linear regression on the data of each cluster and obtain *R*^2^ and corresponding p-value to ascertain the quality of the fit. The *R*^2^ score for five clusters are (0.70, 0.48, 0.60, 0.70 & 0.60) with p-value < 0.05 for all. This implies that, these percentages of variances in 〈CD〉 are explained by mean 〈PD〉 in each cluster. The regression lines on the data for all the clusters are shown with blue solid lines in the plots.

The second observation is related to the ranges of 〈CD〉 and 〈PD〉 for basins within the clusters. These ranges can be read from the Figure. 2 (b) for all five clusters. We observe that both 〈CD〉 and 〈PD〉 ranges are towards higher values ([〈CD〉 >= 0.02] and [〈PD〉 >= 200]) for three clusters (1, 2, 4) out of five. This implies that these basin groups tend to have higher higher mean phylogenetic distance and higher mean cosine distance between the morphological traits of the species within them than the other two. For more clarity, we prepare clusters of randomly selected basins (from all the 2073 HUC8 regions), of equal size to the actual clusters shown in part (a). For all five random clusters, shown in Figure. 2 (c), although we again obtain significant correlation between the 〈CD〉 and 〈PD〉, the ranges of the quantities also cover the lowest values of the respective means, i.e. 〈CD〉 <= 0.02 and 〈PD〉 <= 200, apart from higher values. This is also true for other randomizations (not shown) over basin clusters. The clusters (1, 2, 4) also mostly comprise the species rich eastern region of the US. Hence, on comparison with the random scenario, we conclude that the three basin clusters (1, 2, 4) that have higher number of common species, avoid lower 〈CD〉 and 〈PD〉 values. In other words, species in basins with higher co-occurrences (in eastern part of the US) stick to larger 〈CD〉 and 〈PD〉 ranges, whereas this cannot be strictly said for the western part (3, 5 clusters) of the US. Cluster 5 also has a uniquely high number of basins with high mean morphological distance, which means that a few basins in the west have the most morphologically distinct species.

### B. Primary determinant of species co-occurrence: phylogeny or morphology?

In the previous sub-section, we established that basin level 〈PD〉 and 〈CD〉 are correlated quantities for US freshwater fish species, irrespective of the geographical location of the basin in consideration. In our quest of understanding if and to what degree, do these quantities correlate or explain variances in number of basins of cooccurrences, the already established robust correlation between 〈PD〉 and 〈CD〉 hint that it is safe to expect that either both or neither of these quantities explain variance in chances of species co-occurrence. However, if both these quantities correlate with the species chances of cooccurrence, the question as to which of these quantities is a more fundamental or a stronger determinant of species co-occurrence still stands. In this sub-section our focus is to analyze PD and CD based species networks to ascertain which among these two quantities is a stronger determinant of species co-occurrence. These networks for CD metric for two nearest neighbour choices, NN = 10 and NN = 50 are shown in Figure. 7 in SI, with clusters obtained from the community detection procedure shown in different colours. For the higher number of NN, the networks are organized into smaller number of modules and the clustering is of lower quality. This can be understood from the plot in Figure. 6 (a-b) in SI showing decreasing value of modularity (*Q*) as a function of NN choice for both CD based and PD based networks.

To understand if the species co-occurrences are dictated by their trait dissimilarity, i.e., to test hypothesis *H*_1_ as defined in methods section, we obtain the correlation of 〈CD〉 and NBS, for the CD-based networks for a range of NN values (NN = 1 to NN = 100). These plots are shown in Figure. 3, for NN = 10 (a) and NN = 50 (b) where data is sorted in ascending order of NBS. The horizontal dashed blue line is the mean over 〈CD〉 of all clusters. For each cluster in the CD-based network, we also obtain the 〈PD〉 of species within them. The red dots and the dashed red horizontal line in the same plots shows cluster-wise 〈PD〉 and mean over 〈PD〉 of all the clusters, respectively. From these two plots, it appears that NBS increases with 〈CD〉 with some exceptions. Additionally, we observe that 〈CD〉 and the corresponding 〈PD〉 fluctuate similarly around their mean values (red and blue horizontal lines). To statistically test the dependence of NBS on 〈CD〉, i.e. to test our hypothesis Hi, we obtain *R*^2^ coefficient and p-value with 〈CD〉 as the independent variable and NBS as the dependent variable (see values in Table. I). From this analysis, we find that the dependence is significant (*p* – *value* < 0.05) for NN = 10 network but not for NN = 50 network. On performing similar test with 〈CD〉 as the independent variable and 〈PD〉 as the dependent variable, we again find the dependence is significant (*p* – *value* < 0.05) for NN = 10 network but not for NN = 50 network.

**FIG. 3.**
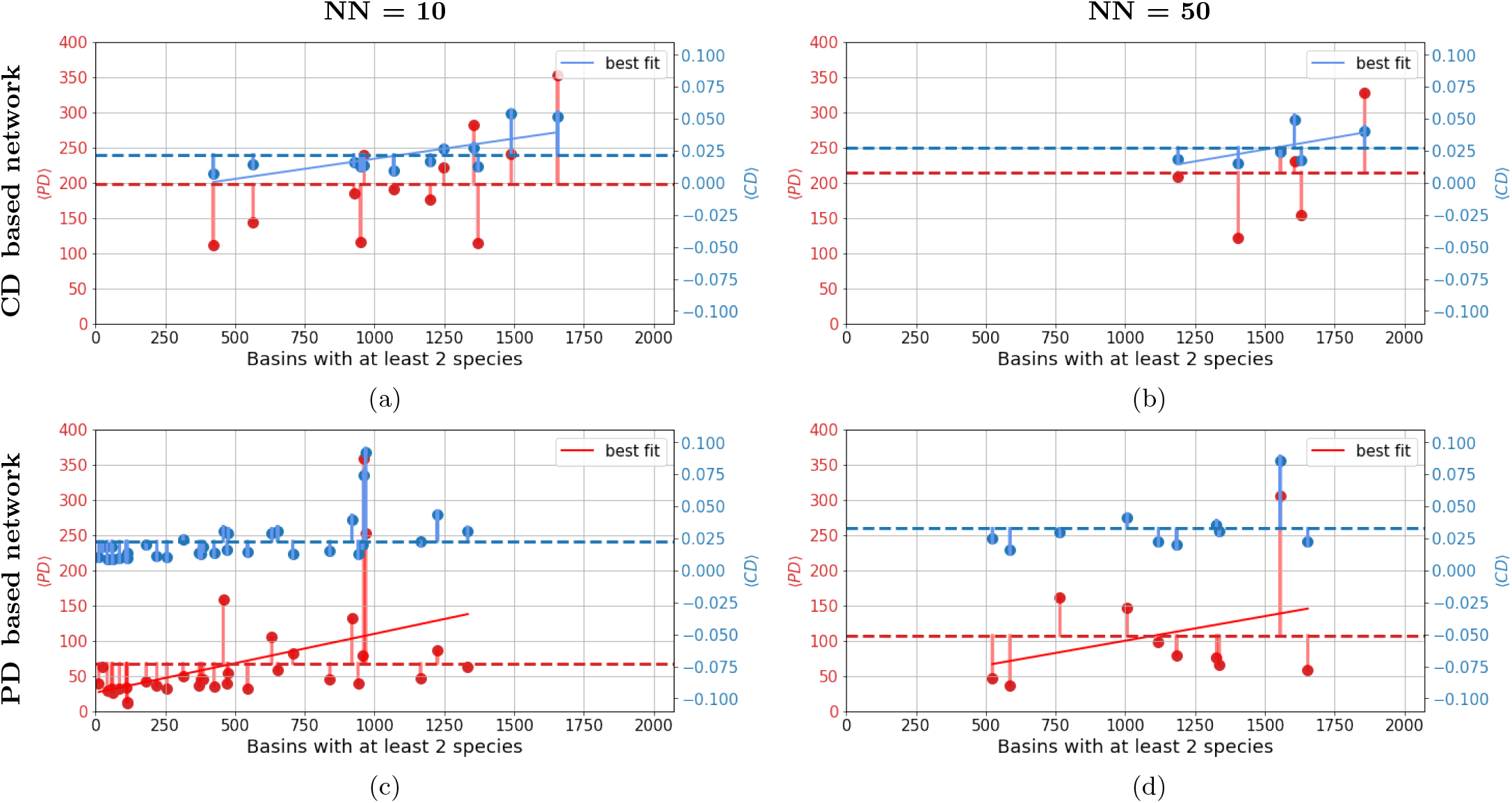
Relationship between cosine distance, phylogenetic distance and number of basins of co-occurrence based on species networks. For the network constructed using cosine distance between species, the figure show relationship between the number of basins having at-least two of the species of the module (along x-axis) and corresponding means 〈CD〉 (right y-axis) and 〈PD〉 (left y-axis) of species within modules, for NN = 10 (a), and NN = 50 (b) networks. Similarly, the bottom row shows corresponding results for species network constructed using PD between species for NN = 10 (c), and NN = 50 (d).

**TABLE 1.** Summary statistics of linear regression models showing relation among three variables: number of basins of co-occurrence of at least two species (NBS), PD and CD. *x* and *y* represent independent and dependent variables, respectively, in the regression analysis. Results for four settings with respect to the metric used of species network construction and the number of nearest neighbours are shown. The *R*^2^ coefficient and the p-values shown here are for a single experiment.

|  | NN = 10 |  |  |  | NN = 50 |  |  |  |
| --- | --- | --- | --- | --- | --- | --- | --- | --- |
| Metric used | y | x | $R^2$ | p-value | y | x | $R^2$ | p-value |
| CD | NBS | $\langle CD \rangle$ | 0.412 | **0.024** | NBS | $\langle CD \rangle$ | 0.256 | 0.305 |
| | $\langle PD \rangle$ | $\langle CD \rangle$ | 0.604 | **0.002** | $\langle PD \rangle$ | $\langle CD \rangle$ | 0.487 | 0.122 |
| PD | NBS | $\langle PD \rangle$ | 0.217 | **0.006** | NBS | $\langle PD \rangle$ | 0.111 | 0.345 |
| | $\langle CD \rangle$ | $\langle PD \rangle$ | 0.790 | **<<0.05** | $\langle CD \rangle$ | $\langle PD \rangle$ | 0.838 | **0.0001** |

To check hypothesis *H*_2_, we do similar analysis with PD-based network, i.e. investigate if 〈PD〉 can explain the variance in NBS. The results are shown in Figure. 3 (c-d) in SI. In this case, we see a clear trend of variation of the quantities among each other for NN = 10 (c) network and for NN = 50 network. To ascertain the significance of observed relations between the three quantities, we obtain linear regression statistics for these networks in terms of R^2^ coefficient and p-value as shown in Table. I for these particular nearest neighbour choices. For the NN = 10 PD-based network we find that the 21.7% of the variance in NBS (R^2^ = 0.217, *p* < 0.05) is significantly explained by 〈PD〉. For NN = 10 CD-based networks we observe significant correlations between NBS and 〈CD〉 (R^2^ = 0.412, *p* < 0.05). The R^2^ values for NN = 50 networks are also presented alongside which tell that correlation between NBS and 〈CD〉 (for CD based network) and correlation between NBS and 〈PD〉 (for PD based network) are not significant. This motivated exploring statistics for a range of NN values for both these network types. Restricting to NN values that lead to higher modularity values of obtained networks and hence result in meaningful modules (see Figure. 6), we chose the range 2 < NN < 100 for both the network types. The statistics for the full range of nearest neighbour choices are presented in plot (a) in Figure. 4, which shows *R*^2^ values, with *p* – *value* < 0.05 shown by filled circles and *p* – *value* ≥ 0.05 shown using hollow circles. All the results in this figure are averages over 100 runs of module identification algorithm for each NN choice, to account for any randomness in module assignment.

**FIG. 4.**
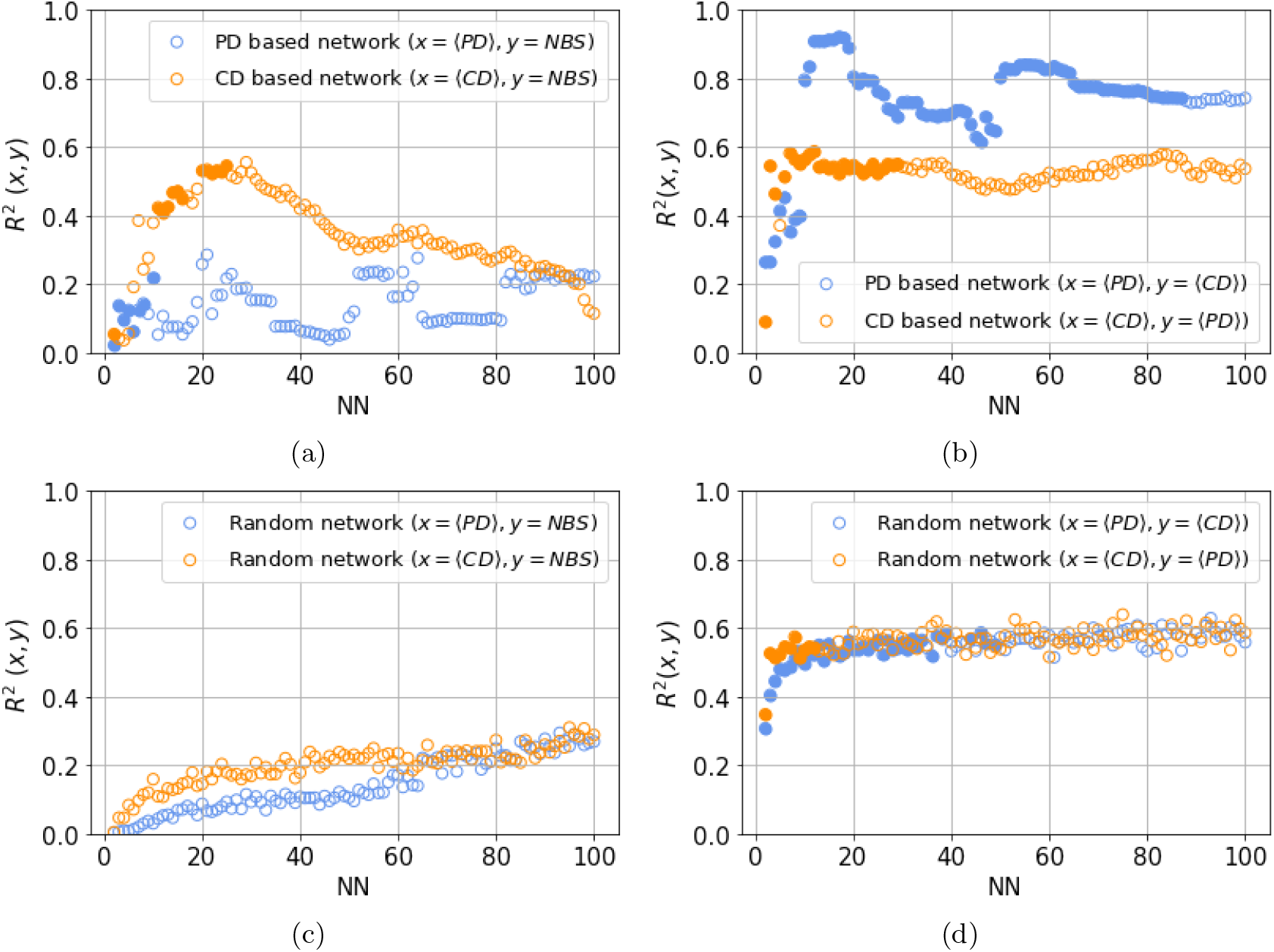
(a-b) Variation of *R*^2^ with NN for CD and PD based networks with *y* are dependent variable and *x* as independent variable. For the CD based network, 〈CD〉 is the independent variable, and for the PD based network 〈PD〉 is the independent variable. The dependent variables in the former case is are NBS and 〈PD〉 and in the latter case they are NBS and 〈CD〉. The filled circles indicate that corresponding *p* – *value* < 0.05, and hollow ones indicate *p* – *value* ≥ 0.05 (c-d) These are the corresponding plots for the random module analysis.

From the full range plots (Figure. 4 (a)), we observe that there exist (almost) distinct ranges of NN values for which the hypotheses *H*_1_ and *H*_2_ can be rejected. For the PD-based networks, this range is 1 ≤ NN ≤ 10, but for the CD-based networks, the range is 10 < NN ≤ 23. Additionally, we observe that for the CD-based network, the significant *R*^2^ values (〈CD〉 explaining variance in NBS) are higher 0.4 < *R*^2^ < 0.6 than the PD-based network where significant *R*^2^ values (〈PD〉 explaining variance in NBS) are 0.0 < *R*^2^ <= 0.2. From the latter observation, we can infer that NBS is more strongly dependent on 〈CD〉 than 〈PD〉. Next in our the full range analysis, we return to correlations between 〈CD〉 and 〈PD〉. For the CD-based network, we explore if 〈CD〉 explains variance in 〈PD〉, and for the PD-based network, we explore how 〈PD〉 explains variance in 〈CD〉. We observe, as shown in plot (b) of the Figure. 4, that there is much longer range of NN values where 〈PD〉 explains variance in 〈CD〉 than the case where 〈CD〉 explains variance in 〈PD〉. This observation reflects that species clusters obtained based on phylogenetic closeness tend to incorporate morphologically similar species for a larger range of nearest neighbours.

So far we have not given any biological meaning to modules we identify, nor do we claim that these are the unique representations of phylogenetically close or morphologically similar species. However, to understand if (indirect) evidence points to the relevance of these modules and to check for the validity of our results, we examine the statistics obtained from random modules of species. To this end, we randomly select species and consider them as modules such that they are of the same sizes as the actual modules for the whole NN range that we are interested in. We observe that neither 〈CD〉 nor 〈PD〉 explains variance in NBS for any of the NN values. On the other hand, both 〈CD〉 and 〈PD〉 are correlated with each other, but for a smaller range of NN than with analysis using the actual networks’ module. The plots for the random module analysis are shown in Figure. 4(c-d). Note that, here too the range of NN for 〈PD〉 significantly explaining variance in 〈CD〉 is longer than the reverse case. Both these observations argue in favour of the modules we identify and the conclusions we draw from their analysis.

## V. DISCUSSION

In this work, we set out to understand how phylogenetic diversity is related to (or explains) morphological diversity of fish species and if morphology and phylogeny govern the species content of ecological fish communities. We use the phylogenetic distances, morphlological traits, and occurrence information of native fish species in the HUC8 regions of the US. Clearly, the dataset we use in our experiments accounts for a large geographical expanse in terms of species occurrence and also accounts for all native fishes in the US. To the best of our knowledge our study is one of the first to utilize continental level fish data for this domain of research.

Traditionally ecologists have relied on using *Phylogenetic Signal of morphological traits* to understand statistical dependency between trait values of species as a consequence of their phylogenetic relation. Although some recent studies have pointed out that the use of phylogenetic signal is not always reliable [8, 9], it is still the state-of-the-art method used in most ecological studies. We present the phylogenetic signal on our dataset in Table II. The values of Bloomberg’s *K* and Pagel’s λ suggest existence of strong phylogenetic signal for native fish species in the US. This means that the morphological distance is statistically explained by phylogentic distance. Our basin network analysis, not only (broadly) confirms this but also shows how this relation varies in different geographical regions across the US, by design. Through a comparison with random basin groups, we find that basins (HUC8 regions) in the species rich East end of the map show stronger statistical dependence between these quantities than the rest of basins. These basins also happen to be those that have *only* higher values of mean phylogenetic distances and higher mean morphological distances given the full range these quantities take for all the basins.

**TABLE II.**
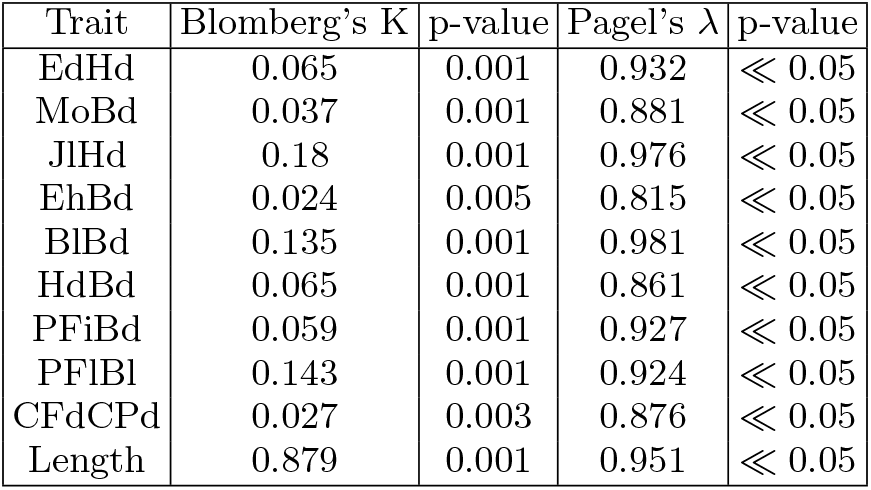
Phylogenetic signal of morphological traits

The CD based and PD based species network analysis reveals that both phylogenetic distance and morphological dissimilarity determine the chances of species co-occurrences, but between these two, morphological dissimilarity between species more strongly determines if they co-occur or not than the phylogenetic distance. These observations are made for specific values of NN for the network types, and there are NN values for which the null-hypotheses could not be rejected. The larger NN values where the latter is true, results in networks with smaller number of modules of large sizes, which are also not very discrete, i.e. there are a increased number of inter-modular edges. These large modules although show similar ranges of variation in 〈CD〉 and 〈PD〉 (as for low NN), the NBS values are restricted to a very small range towards very high values. The smaller number of data points (modules) for higher NN values might have lead to the correlations being not captured with statistical significance. Nevertheless, since meaningful modules pertain to good modularity scores (low NN values), our results still stand firmly where this is true.

For our null-hypotheses tests pertaining to these observations, we constructed networks of species where interactions between species pairs are assigned if phylogenetic and morphological distances are small. As a result, the modules in these networks are determined by and have small phylogenetic and morphological distances between species. For CD based networks, the overall 〈CD〉 values are small, as expected, and similarly for PD based networks the 〈PD〉 values of different clusters are small when compared to the overall ranges CD and PD can take as shown in histograms for CD and PD in Figure. 5 (a-b) in the SI. Although 〈CD〉 and 〈PD〉 values of the clusters are low in the respective networks, they vary; and this allows to investigate how the number basins of co-occurrence (NBS) of species within the clusters vary with 〈CD〉 and 〈PD〉 of the clusters. Hence, on studying of variation of NBS with 〈CD〉 and 〈PD〉 we could infer statistical dependence between them. The methodology used in this work, involves nearest neighbour species networks that are constructed based directly on the CD and PD values and the network modules are simple groups of species that share most neighbours among themselves. The analyses based on these groups to derive the dependencies between the quantities (morphological dissimilarity, phylogenetic distance and chances of species co-occurrence) allows for a complex understanding of their relations as compared to looking at variations of overall mean values of these quantities against each other.

**FIG. 5.**
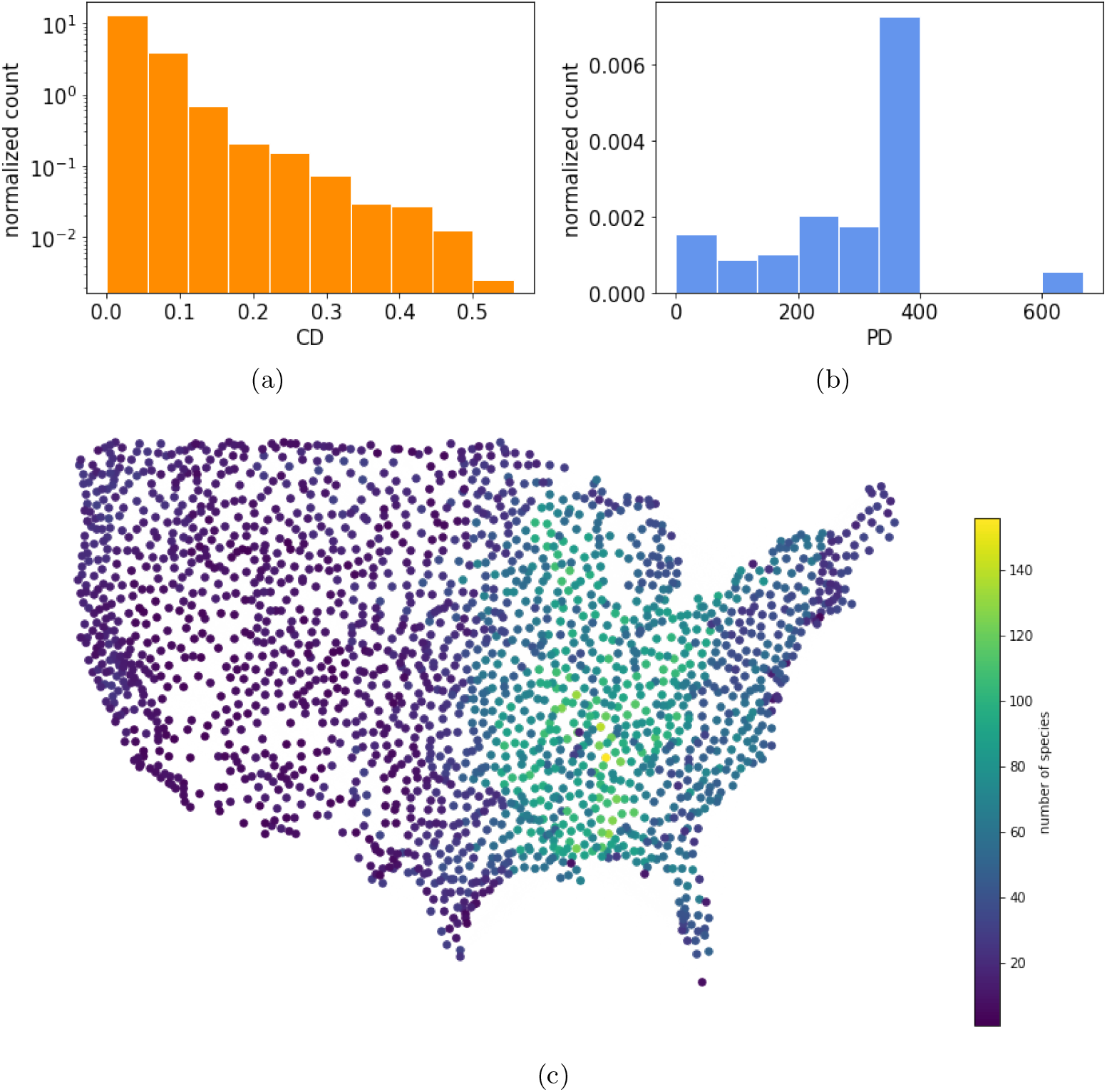
(a-b) Distribution of Phylogenetic distances and Cosine distances between fish species (c) Map of number of species in each basin: The dots indicate the centroids of HUC8 regions and colours the number species present in the region. The range of species numbers is shown in the color map along side.

The two main observations from our study, are 1) that phylogenetic distance explains the variance in morphological dissimilarity of fish species, and 2) that morphological relationships among the species govern chances of species co-occurrence more strongly than their phylogenetic relationship. The first observation points to the important role that evolutionary history of species play in their form and function; and the second to the role of species morphology in their probability of co-occurrence.

For the species networks, we limit to NN values in the range NN = 1 and NN = 100. As we include more nearest neighbours in the network, more distant species are allowed to form connections. This in turn results in increasing inter-module edges in the module structure, which is reflected in the decreasing modularity score of the network. Hence, the strength of division of a network into meaningful modules identified by groups comprising the metric-wise closest species, is weakened with increasing NN choices. Since our analyses is based on identifying groups of phylogenetically and morphologically similar species, and then testing the hypotheses about correlation with number of basins of co-occurrence, it is important that the identified groups are indeed metric-wise similar/ closer species. To ascertain this condition, we make use of the modularity value (*Q*), and restrict to NN values with good modularity scores. For the CD based network the *Q* value drops steeply with NN, whereas for the PD based network the decline is gentler. See Figure. 4 (a-b) in SI where *Q*_NN=10,CD_ = 0.7 and *Q*_NN=50,CD_ = 0.55 for CD-based networks and Q_NN=10,PD_ = 0.9 and *Q*_NN=50,PD_ = 0.75 for PD based networks. Apart from modularity score, the number of clusters decreases with increasing NN, due to merger of the modules (obtained at smaller NN with each other). For example, for CD-based networks for NN = 10, the number of clusters was 11, but for NN = 50, the number of clusters was just 6 (see Figure. 7). Similar situation also occurs for the PD based networks.

**FIG. 6.**
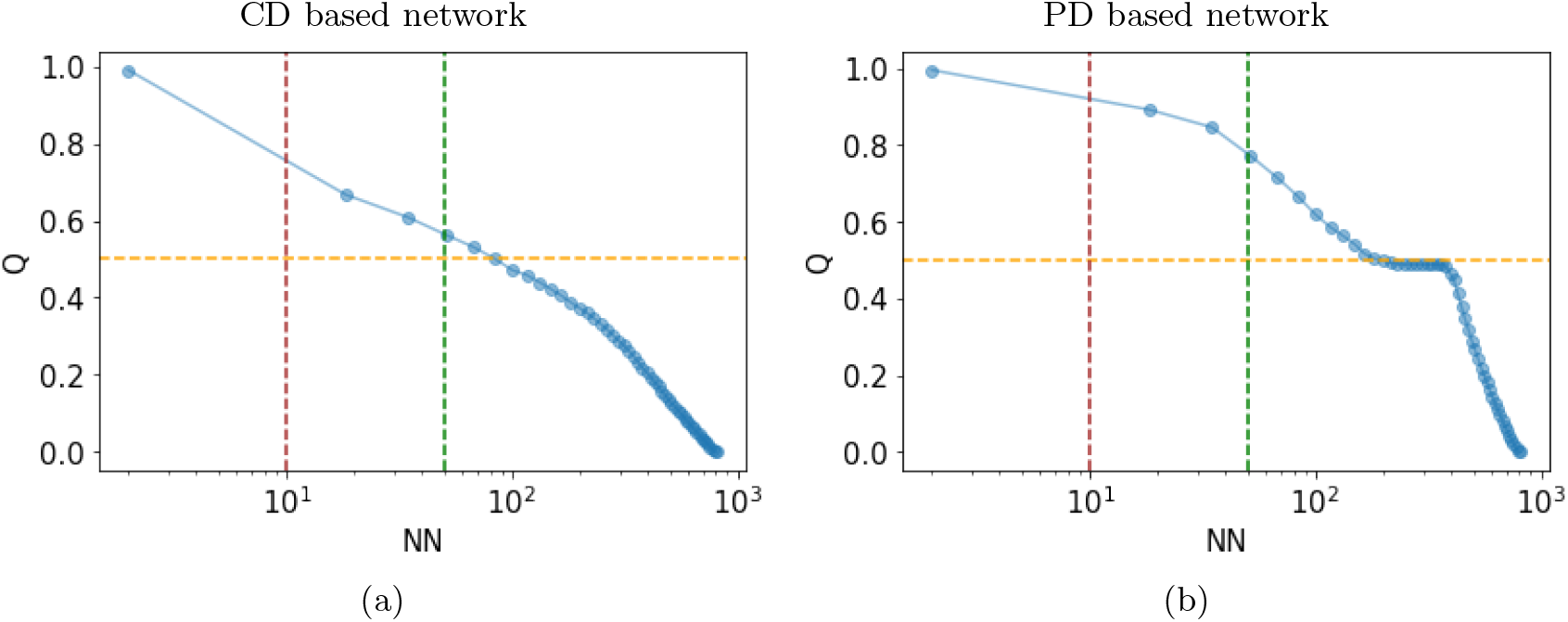
(a-b) Variation of network modularity with number of nearest neighbours. The vertical line mark NN = 10 and NN = 50, and horizontal line marks *Q* = 0.5 for (a) CD based network, and (b) PD based network.

**FIG. 7.**
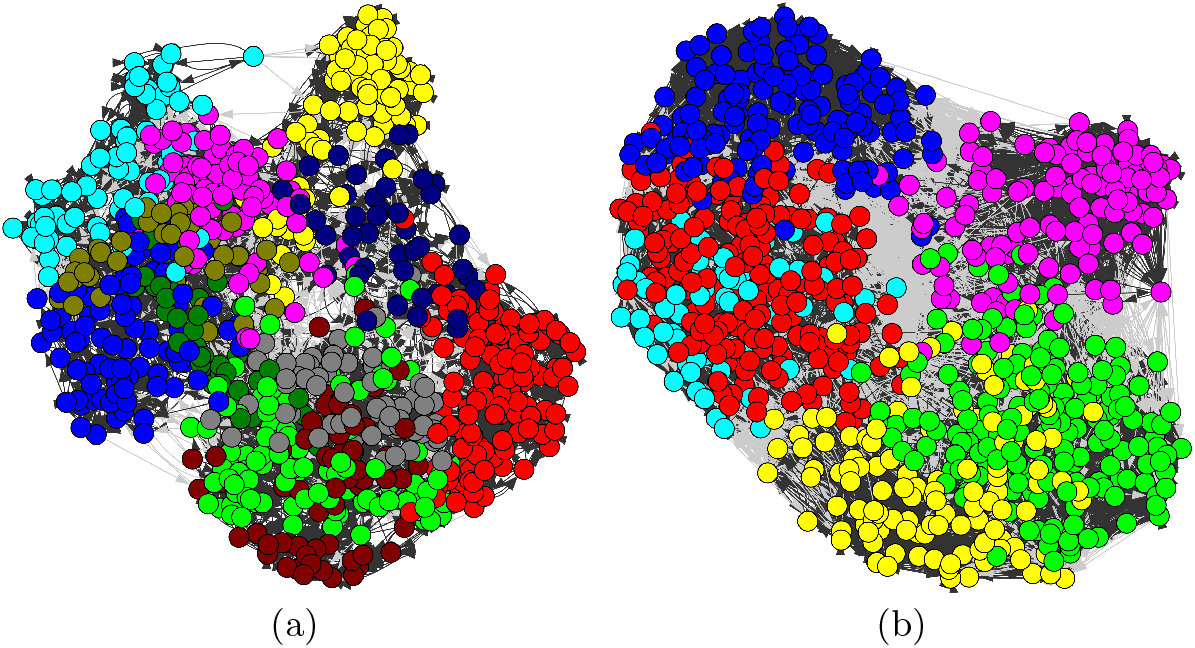
Nearest neighbor networks of species constructed based on Cosine distances between them. (a) The network, where each node is connected to its 10 nearest neighbors, shows 11 different modules or network communities. (b) The network where each node is connected to its 50 nearest neighbors, shows 6 modules. The colours identify different network modules. In both of these networks the directed edges within module are shown in black colour and inter-modular edges are shown with grey colour.

To estimate extent of co-occurrence (in species network analyses), we used “basins with at least two of the species in a module” (NBS) as the measure. So for a basin to qualify as a *co-occurrence basin* for that module of species, it should have at least two of the species in the module, and NBS is the total number of such basins. We did not choose higher percentage of species in the module since there are basins with a very small number species, and this can result in NBS to be zero especially for smaller sized modules. Therefore to avoid the trivial values of NBS being zero and still retaining the definition of co-occurrence, we stick with the bare minimum condition of at least two of the species in the module. We also checked that the results do not qualitatively differ when cooccurrence of a higher percentage of species in the module are used as a criteria,

## VI. CONCLUSION

This work proposes a novel framework for identifying correlations between Phylogeny, morphology, and the co-occurrence of fish species. The complex networks framework, wherein connections are determined through nearest-neighbours, not only allows modeling local node-specific interactions as edges, but these local interactions can cause the overall structure of the network to be comprised of clustered group of species (revealed through module identification) which are functionally meaningful. Through our module-level analyses of species-species networks based on phylogenetic and morphological distances, and the basin network based on number co-occuring species, we uncover the statistical dependence also confirmed by traditional measures like *Phylogenetic signal of morphological traits*, between fish phylogeny and morphology. Additionally, using this framework, we determine which of the two quantities, fish phylogeny and fish morphology, is a stronger determinant of their co-occurrences. We can extract a few take-home messages from our analysis. Firstly, phylogenetic distance (closeness) in fish species explains morphological distance (closeness) among species. Secondly, basins with a higher species richness have higher mean phylogenetic distances and higher mean morphological distance than basins with smaller richnesses. Thirdly, although both phylogentic distance and trait dissimilarity are significant determinants of number of basins of co-occurrence, morphological distance explains greater degree of variance in number of basins of co-occurrence than phylogentic distance.

## VII. FUNDING STATEMENT

This work was partially funded by the Center of Advanced Systems Understanding (CASUS), which is financed by Germany’s Federal Ministry of Education and Research (BMBF) and by the Saxon Ministry for Science, Culture, and Tourism (SMWK) with tax funds on the basis of the budget approved by the Saxon State Parliament. A.R is supported by the research program of the Netherlands Organisation for Scientific Research (NWO).

## VIII. SUPPLEMENTARY INFORMATION

## Notes

### Competing Interest Statement

The authors have declared no competing interest.

## REFERENCES

[1] S. Kelly, R. Grenyer, and R. W. Scotland, Diversity and Distributions 20, 600 (2014).

[2] D. Mouillot, D. R. Bellwood, C. Baraloto, J. Chave, R. Galzin, M. Harmelin-Vivien, M. Kulbicki, S. Lavergne, S. Lavorel, N. Mouquet, et al., PLoS biology 11, e1001569 (2013).

[3] D. Mouillot, S. Villéger, V. Parravicini, M. Kulbicki, J. E. Arias-González, M. Bender, P. Chabanet, S. R. Floeter, A. Friedlander, L. Vigliola, et al., Proceedings of the National Academy of Sciences 111, 13757 (2014).

[4] D. P. Faith, Biological conservation 61, 1 (1992).

[5] M. Winter, V. Devictor, and O. Schweiger, Trends in ecology & evolution 28, 199 (2013).

[6] F. Mazel, A. O. Mooers, G. V. D. Riva, and M. W. Pennell, Systematic Biology 66, 1019 (2017).

[7] S. P. Blomberg, T. Garland Jr, and A. R. Ives, Evolution 57, 717 (2003).

[8] N. Mouquet, V. Devictor, C. N. Meynard, F. Munoz, L.-F. Bersier, J. Chave, P. Couteron, A. Dalecky, C. Fontaine, D. Gravel, et al., Biological reviews 87, 769 (2012).

[9] L. J. Revell, L. J. Harmon, and D. C. Collar, Systematic biology 57, 591 (2008).

[10] M. R. Winston, The American Naturalist 145, 527 (1995).

[11] B. R. Krasnov, M. A. Fortuna, D. Mouillot, I. S. Khokhlova, G. I. Shenbrot, and R. Poulin, The American Naturalist 179, 501 (2012).

[12] C. Baraloto, O. J. Hardy, C. T. Paine, K. G. Dexter, C. Cruaud, L. T. Dunning, M.-A. Gonzalez, J.-F. Molino, D. Sabatier, V. Savolainen, et al., Journal of ecology 100, 690 (2012).

[13] J. Bascompte, Basic and Applied Ecology 8, 485 (2007).

[14] L. Götzenberger, F. de Bello, K. A. Bråthen, J. Davison, A. Dubuis, A. Guisan, J. Lepš, R. Lindborg, M. Moora, M. Pörtel, et al., Biological reviews 87, 111 (2012).

[15] J. M. Levine, J. Bascompte, P. B. Adler, and S. Allesina, Nature 546, 56 (2017).

[16] M. J. Spasojevic and K. N. Suding, Journal of Ecology 100, 652 (2012).

[17] P. R. Peres-Neto, Oecologia 140, 352 (2004).

[18] C. O. Webb, D. D. Ackerly, M. A. McPeek, and M. J. Donoghue, Annual review of ecology and systematics 33, 475(2002).

[19] D. L. Galat, C. R. Berry, W. M. Gardner, J. C. Hendrickson, G. E. Mestl, G. J. Power, C. Stone, and M. R. Winston, (2005), https://digitalcommons.unl.edu/nebgamestaff/54/.

[20] G. Su, S. Villéger, and S. Brosse, Global Ecology and Biogeography 28, 211 (2019).

[21] D. L. Rabosky, J. Chang, P. F. Cowman, L. Sallan, M. Friedman, K. Kaschner, C. Garilao, T. J. Near, M. Coll, M. E. Alfaro, et al., Nature 559, 392 (2018).

[22] R. C. Bastos, L. S. Brasil, J. M. B. Oliveira-Junior, F. G. Carvalho, G. D. Lennox, J. Barlow, and L. Juen, Ecological Indicators 122, 107257 (2021).

[23] E. L. Rezende, P. Jordano, and J. Bascompte, Oikos 116, 1919 (2007).

[24] C. Fontaine and E. Thébault, Population Ecology 57, 29 (2015).

[25] D. J. McGarvey and J. A. Veech, Plos one 13, e0208720 (2018).

[26] M. Layeghifard, P. R. Peres-Neto, and V. Makarenkov, Molecular phylogenetics and evolution 64, 190 (2012).

[27] S. Brosse, N. Charpin, G. Su, A. Toussaint, G. A. Herrera-R, P. A. Tedesco, and S. Villéger, Global Ecology and Biogeography 30, 2330 (2021).

[28] G. Su, M. Logez, J. Xu, S. Tao, S. Villéger, and S. Brosse, Science 371, 835 (2021).

[29] R. Froese, H. Winker, G. Coro, N. Demirel, A. C. Tsikliras, D. Dimarchopoulou, G. Scarcella, W. N. Probst, M. Dureuil, and D. Pauly, ICES Journal of Marine Science 75, 2004 (2018).

[30] C. Penone, A. D. Davidson, K. T. Shoemaker, M. Di Marco, C. Rondinini, T. M. Brooks, B. E. Young, C. H. Graham, and G. C. Costa, Methods in Ecology and Evolution 5, 961 (2014).

[31] D. J. Stekhoven and P. Bühlmann, Bioinformatics 28, 112 (2012).

[32] G. Su, A. Mertel, S. Brosse, and J. M. Calabrese, bioRxiv (2022), https://doi.org/10.1101/2022.03.04.481515.

[33] E. Paradis, S. Blomberg, B. Bolker, J. Brown, J. Claude, H. S. Cuong, R. Desper, and G. Didier, Analyses of phylogenetics and evolution, version 2, 47 (2019).

[34] G. Sidorov, A. Gelbukh, H. Gómez-Adorno, and D. Pinto, Computación y Sistemas 18, 491 (2014).

[35] N. Reyes, R. Connor, N. Kriege, D. Kazempour, I. Bartolini, E. Schubert, and J.-J. Chen, Similarity Search and Applications, Vol. 13058 (Springer Nature, 2021).

[36] P. Xia, L. Zhang, and F. Li, Information Sciences 307, 39 (2015).

[37] M. E. Newman and G. Reinert, Physical review letters 117, 078301 (2016).

[38] V. D. Blondel, J.-L. Guillaume, R. Lambiotte, and E. Lefebvre, Journal of statistical mechanics: theory and experiment 2008, P10008 (2008).

[39] M. E. Newman, Physical Review E 94, 052315 (2016).

[40] V. A. Traag, L. Waltman, and N. J. Van Eck, Scientific reports 9, 1 (2019).

